# Computationally Guided Design of Two Novel Phosphodiesterase-4 Inhibitors for Treatment of Plaque Psoriasis

**DOI:** 10.64898/2026.10.20.760516

**Authors:** Annie Liu, Emma R. M. James, Justine B. Siegel

## Abstract

Plaque psoriasis is a chronic inflammatory skin disease characterized by itchy, scaly skin. Although there is currently no cure for this disease, it can be managed with medication, such as phosphodiesterase-4 (PDE4) inhibitors. Current PDE4 inhibitors are limited and show adverse effects. With the goal to develop better PDE4 inhibitors for the treatment of plaque psoriasis, two novel drug candidates were designed with computational studies, and observed to have better ADMET properties compared to Roflumilast. A homology analysis was performed to determine the best model organism for preclinical studies of the novel candidates.

## INTRODUCTION

Psoriasis is a chronic, inflammatory skin condition. Plaque psoriasis is the most common manifestation of psoriasis. Approximately 80-90% of psoriasis cases are classified as plaque psoriasis with the disease affecting an estimated 83-250 million people worldwide. Psoriasis and plaque psoriasis are both characterized by skin irritation, with plaque psoriasis having an emphasis on silvery, scaly patches on the skin, commonly affecting the elbows, back, knees, and scalp.^1,2,3^ This condition can significantly reduce quality of life and is associated with complications such as psoriatic arthritis, cardiovascular disease, metabolic syndrome, and mental disorders.^1,3^ Due to limited treatment options and no current cure, managing plaque psoriasis and its consequences remains a challenge.

With the uprise of new generation of novel drugs, effective treatment options include phosphodiesterase-4 (PDE4) inhibition which has shown promise in managing the symptoms of plaque psoriasis.^4,5^ PDE4 regulates immune responses in the body by maintaining levels of cyclic adenosine monophosphate (cAMP), a signaling molecule involved in immune cell communication.^4,5^ PDE4 catalyzes the hydrolysis of cAMP through cleavage of phosphodiester bonds.^4^ Inhibiting PDE4 increases cAMP levels, which leads to anti-inflammatory outcomes.^5^ Previous studies have shown that the PDE4B subfamily is selectively involved in specific inflammatory responses, making it a key target in the development of PDE4-specific anti-inflammatory drugs.^5,11^ However, due to the similarities of all subfamilies of PDE4, unintended inhibition of the PDE4D subfamily in the brain can cause adverse side effects such as nausea and vomiting, highlighting importance of subfamily selectively in novel drug design.^5^

Previously known PDE4 inhibitors, for the treatment of plaque psoriasis, will serve as the basis for the design and evaluation of new novel PDE4 inhibitors. These previously known inhibitors include Roflumilast, Crisaborole, Orismilast, and Mufemilast (Figure 1).^6,7,8,9,15^ Among these, only Roflumilast is approved by the FDA for topical use in plaque psoriasis.^6,15^ Crisaborole, approved for atopic dermatitis, has also demonstrated potential for topical plaque psoriasis treatment.^7,10^ Orismilast and Mufemilast are oral treatments currently in clinical trials; while not yet FDA approved, both have shown encouraging results for plaque psoriasis.^8,9^ These will serve as the established known PDE4 inhibitors in the PDE4 inhibitor drug development for plaque psoriasis.

**Figure 1.**
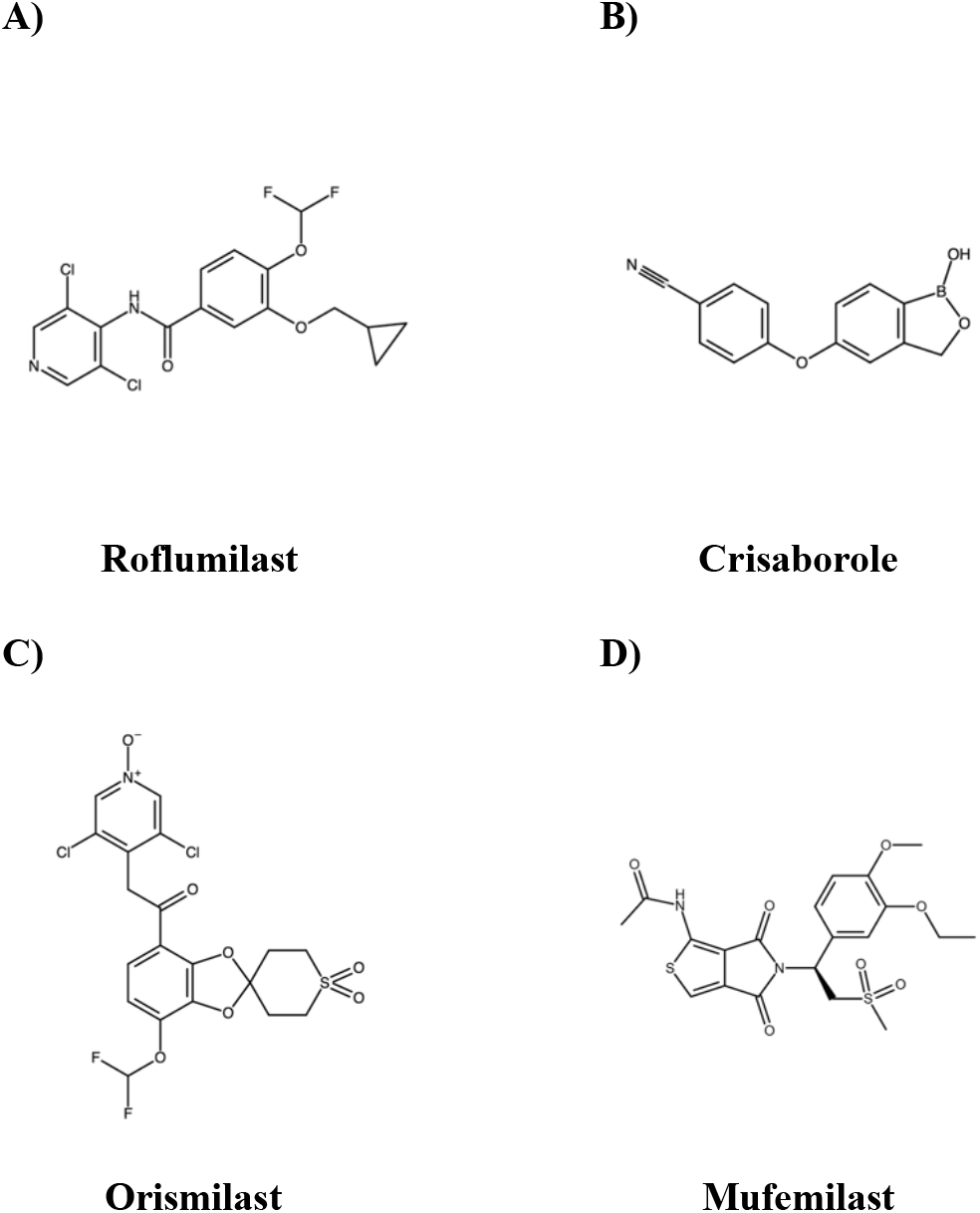
Structures of previously known PDE4 inhibitors: A) Roflumilast, B) Crisaborole, C) Orismilast, D) Mufemilast

The safety and side effects of these drugs vary depending on their route of administration, which is an important consideration for clinical use. For the most part, oral drug treatments are preferable and abide by Lipinski’s Rules of 5 to determine good oral bioavailability. Common adverse effects of Roflumilast, Orismilast, and Mufemilast include gastrointestinal discomfort, diarrhea, headache, nausea, and vomiting.^8,9^ Although, these symptoms are generally milder with topical Roflumilast compared to oral administration.^6^ In contrast, topical Crisaborole has not been associated with significant application site irritation or other notable adverse effects.^7^ Gastrointestinal side effects are most commonly reported with oral Orismilast and Mufemilast, likely due to the fact that oral treatments have to go through the gastrointestinal tract.^8,9^ A fundamental understanding of the safety and side effects is essential for designing novel drugs and improving patient outcomes in plaque psoriasis.

However, most current PDE4 inhibitors lack high subfamily specificity, which increases the risk of adverse effects. Only a few have received FDA approval for plaque psoriasis treatment, providing the motivation for the need of more selective and effective drug treatments for this disease. This limitation has made the development of new, more specific PDE4 inhibitors an active area of research.

A detailed understanding of the PDE4 active site is essential for designing and developing novel drugs, especially through computational studies. The active site consists of three main pockets: the metal binding pocket (M pocket), the glutamine hydrogen bonding pocket (Q pocket), and a solvent filled pocket (S pocket).^5^ The Q pocket plays a crucial role in ligand binding, as cAMP and other high-affinity ligands form hydrogen bonds with an invariant glutamine in this Q pocket.^5^ This glutamine can form either a single or bidentate hydrogen bond, influencing ligand binding affinity.^5^ Previous research studies have identified dialkoxyphenyl (Figure 2) as the essential pharmacophore for PDE4 binding, as it maximizes hydrogen bond formation with glutamine.^5,12^ Additionally, the two metal ions in the M pocket, zinc and magnesium, enhance ligand binding through metal ion interactions.^5^ Both glutamine hydrogen bonding and metal ion interactions are the central hypothesis in the design of new PDE4 inhibition.

**Figure 2.**
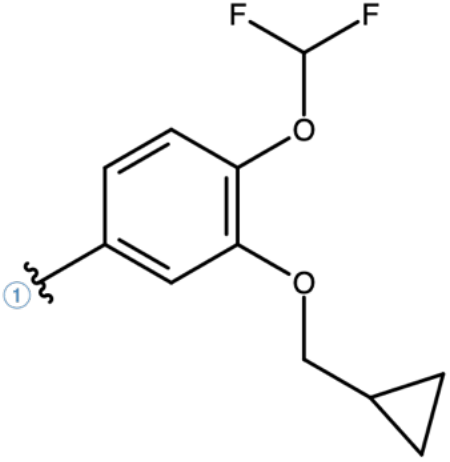
Structure of dialkoxyphenyl, pharmacophore of PDE4 inhibitor drugs.

In effort to find treatment for plaque psoriasis, this study aimed to design and evaluate two new PDE4 inhibitor drug candidates. Using computational methods and following the proposed pharmacophore of dialkoxyphenyl and active site interactions, Candidate 1 was developed through computational modeling and Candidate 2 was designed using chemical intuition. The safety and efficacy of both candidates were evaluated by comparing their docking scores, absorption, distribution, metabolism, excretion, toxicity (ADMET) properties, and active site interactions with those of previously established PDE4 inhibitors. The two candidates show promising potential to be novel drug inhibitors of PDE4. To support future preclinical studies, a homology analysis was performed to propose which organisms are suitable for evaluation of novel drug candidates. Additional homology analysis of pathogenic and beneficial microbes was performed to assess potential off-target effects from other organisms bearing homologous proteins.

## METHODS

The crystalline structure of phosphodiesterase-4B enzyme (PDB ID: 1XMU)^14^ was obtained from the RCSB Protein Data Bank. Make Receptor was used to create the active site of PDE4B after retrieval from PDB.^17^ GaussView was used to build the four inhibitor molecules, followed by Gaussian09 to optimize the molecules.^18,19^ Visualization of the drug and active site and its interactions were done through VIDA and Pymol.^20,21^

OpenEye OMEGA was used to generate conformer libraries of the drugs after optimization.^22^ OpenEye FRED was used to dock the conformer libraries of the drugs into the active site.^23^ OpenEye FILTER was used to screen for ADMET and structural properties.^24^ vRocs was used to screen for electrostatic properties and structural similarity to cAMP.^25^ This led to generation of docking scores, ADMET values, and TanimotoCombos, all of which were considered when determining the lead target for the design of two novel drugs. The lead target was determined through evaluation of docking score, ADMET values, and structural similarities, and which shows most potential for improvement in PDE4 inhibition. Then, the selected lead target was loaded up in vBrood to generate bioisosteric analogs to create Candidate 1.^26^ Candidate 2 was designed from chemical intuition. Docking scores were obtained once again for both candidates, and final screenings followed through OpenEye FILTER to obtain ADMET properties.

Lastly, homology analyses were performed using BLAST to search for the homologous proteins, and Jalview to analyze the sequence alignment of homologous proteins of the organisms.^27,28^

## RESULTS AND DISCUSSION

### Evaluation of known PDE4 inhibitors

To identify the lead target for designing two novel drugs, the four known PDE4 inhibitors were first built and optimized. Computational studies evaluated their docking scores, ADMET properties, and active site interactions to establish preliminary guidelines for the design of new PDE4 inhibitors. The four known inhibitors (Roflumilast, Crisaborole, Orismilast, and Mufemilast) were then compared for safety and efficacy. All computational modeling used the active site of PDE4B, as inhibition of this subfamily of PDE4 enzyme is considered ideal for anti-inflammatory effects in the treatment of plaque psoriasis.^5^

Before performing computational tests of the four known PDE4 inhibitors, an initial control was performed by docking cAMP into PDE4B active site (PDB: 1XMU)^14^. Prior to docking, the molecular structure of cAMP was obtained through PDB: 2PW3,^13^ which was then optimized and conformer libraries were generated. The active site model showed that the glutamine in the Q pocket hydrogen bonds with the adenine base at 1.8 Å and that the cyclic phosphate group interacts with the metal ions, magnesium (MG) and zinc (ZN) at 2.4 Å and 2.7 Å, respectively (Figure 3A). In addition, the adenine base was observed to have a displaced Pi-Pi stacking interaction with F446 at 3.7 Å (Figure 3B).

**Figure 3A.**
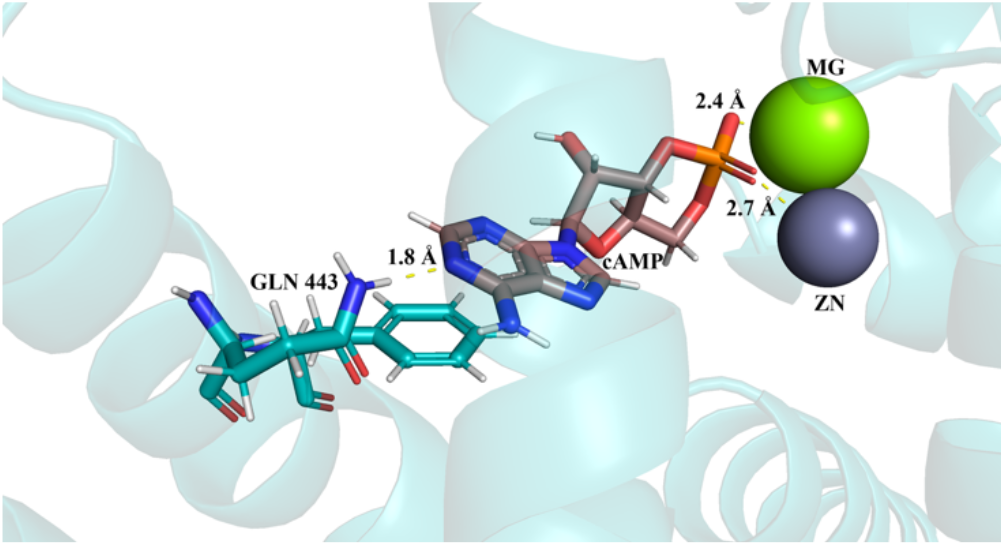
PDE4B active site with predicted cAMP binding: cAMP adenine base (brown) hydrogen bonding with Q443 (cyan), and pyridine interacting with metal ions (green and grey).

Following the initial control docking, computational simulations were conducted to determine which of the four known PDE4 inhibitors would serve as the lead target for novel drug design. All four inhibitor molecules were constructed and optimized and conformer libraries of Roflumilast, Crisaborole, and Mufemilast were generated. However, Orismilast could not be properly evaluated for binding affinity due to a parameter failure in OMEGA, which prevented the generation of a conformer library and docking score calculation.^22^

**Table 1.** Docking scores and ADMET properties of existing inhibitors of PDE4.

| Name | Docking Score | Molecular Weight (g/mol) | H-bond Acceptors | H-bond Donors | XLogP |
| --- | --- | --- | --- | --- | --- |
| Roflumilast | -11.50 | 403.21 | 5 | 1 | 4.23 |
| Crisaborole | -8.82 | 251.05 | 4 | 1 | 2.28 |
| Mufemilast | -12.15 | 465.52 | 9 | 1 | 3.14 |

**Figure 4A.**
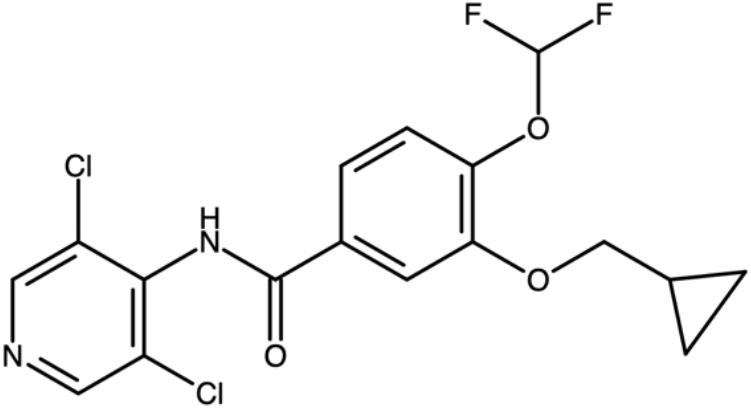
2D structure of Roflumilast.

**Figure 4B.**
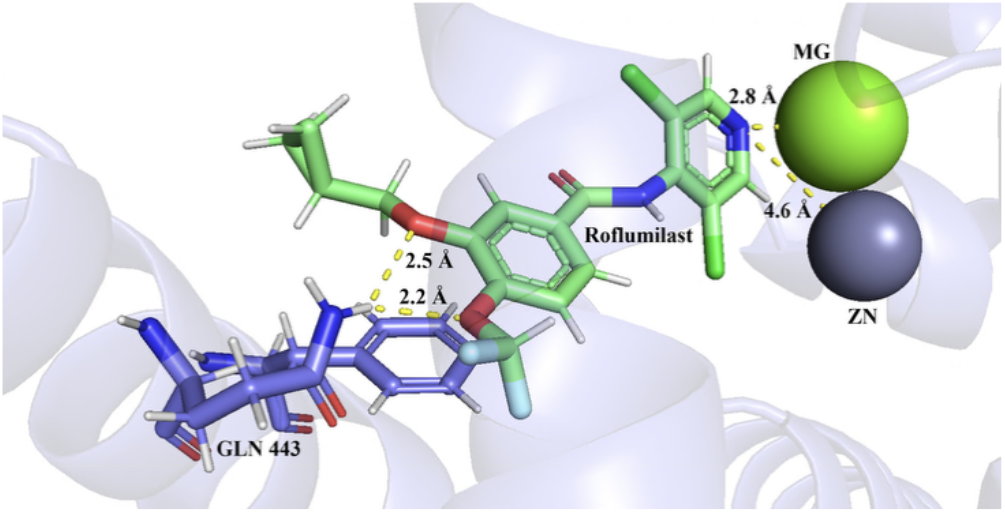
Roflumilast (mint green) bidentate hydrogen bonding with Q443 (blue), and pyridine interacting with metal ions (green and grey).

**Figure 4C.**
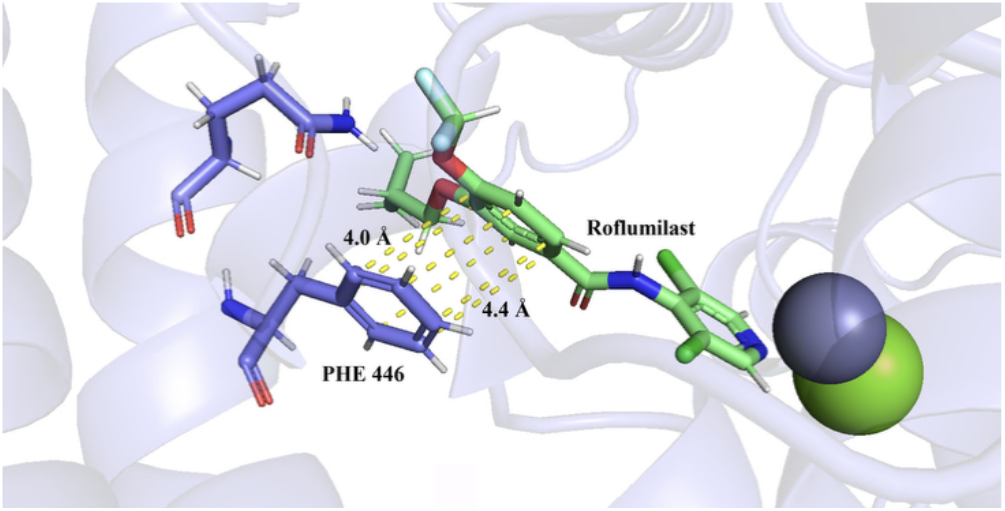
Dialkoxyphenyl (mint green) Pi-Pi stacking with F446 (blue).

The remaining three structures were then docked, screened through FILTER for AMDET properties, and screened through vROCS for structural similarities.^24,25^ Mufemilast had the highest docking score at −12.15, followed by Roflumilast with a docking score at −11.50, while Crisaborole had the lowest docking score at −8.82. Ideal drug treatments are administered orally. Lipinski’s Rules of 5 sets a guideline for if a drug is readily orally bioavailable provided its structural properties. All three inhibitors had favorable predicted bioavailability scores with ABS > 0.55. Crisaborole’s low molecular weight and XLogP, a measure of lipophilicity, suggest it has good orally bioavailable and passes Lipinski’s Rules of 5. However, despite its promising ADMET properties and most overall structural overlap to cAMP, Crisaborole does not contain the essential pharmacophore for inhibition of PDE4B described in previous studies.^5^ Therefore, both Orimilast and Crisaborole were eliminated as potential lead targets.

Of the two remaining known inhibitors, Roflumilast had the lowest docking score as well as the lowest molecular weight. However, its XLogP was the highest at 4.23, making it unfavorable for oral drug treatment. This is consistent with the available Roflumilast treatments limited to topical creams and foams. Both Roflumilast and Mufemilast have the necessary pharmacophore for PDE4B inhibition. Additionally, Roflumilast had the lowest number of hydrogen bond acceptors and the second largest vROCS structural overlap to cAMP, which makes Roflumilast a better target for improvement while maintaining strong binding affinity to the active site of PDE4B. Given pharmacophore presence, structural similarities to cAMP, higher XLogP, and energetically favorable docking score, Roflumilast is the best lead target for designing novel drug candidates, as there is most room for improvement in binding affinity while pertaining to the active site hypothesis. While Mufemilast shows great potential to be effective for inhibiting PDE4B with strong docking score and low XLogP value, Roflumilast shows potential to be exploited to create novel drug candidates that could be administered orally.

To further characterize ligand-enzyme binding interactions of Roflumilast in effort to support design of novel drug candidates, the active site model with Roflumilast was visualized in VIDA.^20^ The dialkoxyphenyl group was observed to form strong bidentate hydrogen bonds with Q443 at 2.5Å and 2.2Å (Figure 4B). In addition, weaker displaced Pi-Pi stacking interactions were observed between the dialkoxyphenyl group and F446 at about 4.0Å (Figure 4C). Additionally, the dichloro-pyridine moiety was predicted to be slightly coordinated with the metal ion pocket of the active site, which is supported by the cAMP control experiment as an effective interaction for strong binding affinity (Figure 4B).

**Figure 3B.**
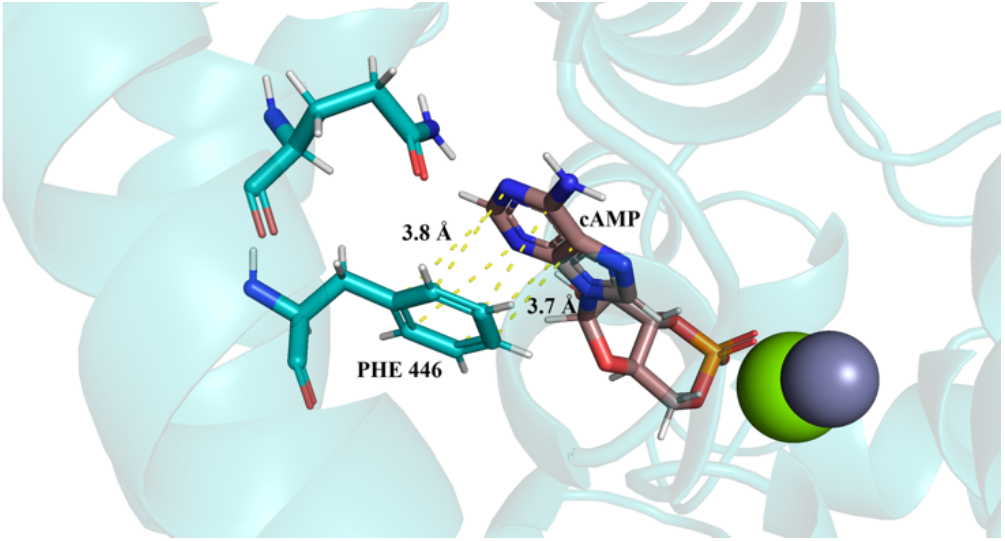
PDE4B active site with predicted cAMP binding: cAMP adenine base (brown) displaced Pi-Pi stacking with F446 (cyan).

Based on these detailed computational analyses, Roflumilast was ultimately selected as the lead target for the design of novel drug candidates. The observed active site interactions of Roflumilast and its molecular structure will serve as a basis for new designs through computational software and chemical intuition, as well as docking scores and ADMET properties.

### Computationally Driven Design of Candidate 1

Using Roflumilast as the lead target, two novel drug candidates will be designed with improvement in efficacy based on docking scores and ADMET properties. Final candidates were screened through CAS SciFinder to ensure originality of the drug molecule.^29^

The first design, Candidate 1 (Figure 5A), was designed from bioisosteric analogs of Roflumilast, computed by vBrood.26 Analysis of the observed Roflumilast and original cAMP substrate binding suggested that the dichloropyridine ring could be further modified to improve upon and optimize interactions with the metal ion coordination pocket of PDE4B. The analysis of the docking scores and structural changes of the generated analogs suggests that the binding affinity is highest when one of chloride substituents on the benzene ring was modified to a methyl group, and the other chloride substituent was modified to a fluorine. The pyridine moiety was retained, but the positions of the fluorine, main chain, and methyl substituents were changed to the ortho-, meta-, and para-position, respectively. This design had the most favorable interaction with the metal ion pocket amongst the bioisosteric analogs generated by vBrood.

Candidate 1 had a docking score of −12.61, which is a 9.7% improvement over Roflumilast. The improvement is largely due to the removal of both chloride atoms, which were replaced with a fluorine and a methyl group, and the repositioning of the substituents on the pyridine ring. Fluorine is more electronegative, and as a result contributes to being a stronger electron withdrawing group. When positioned ortho to the nitrogen and paired with the methyl group in para, the electron density shifts towards the nitrogen and fluorine side of the pyridine ring, increasing the negative charge of the surrounding electron density cloud on the pyridine ring. This negative electron density can interact more strongly with the positively charged metal ions in the PDE4B active site. Importantly, Candidate 1 preserves the dialkoxyphenyl pharmacophore, maintaining its observed bidentate hydrogen bonds with Q443 at 2.3 Å for both hydrogen bonds, and displaced Pi-Pi stacking interaction with F446 at about 4.0 Å, thus maintaining the hypothesis for PDE4B binding (Figure 5A).

It is also important to look at ADMET properties of Candidate 1. Thus, following active site simulation, ADMET properties were generated to assess its overall safety and efficacy. Candidate 1 was predicted to have no violations of Lipinski’s Rules and no detected toxicophores. Additionally, the molecular weight decreased to 366.33 g/mol and XLogP was reduced to 3.81 compared to Roflumilast, both favorable values for oral bioavailability. Although the number of hydrogen bond acceptors and donors did not change, the improved binding interactions described previously contributed to a stronger overall binding affinity, as shown by the improved docking score. The improved docking score and active site interactions show that Candidate 1 is a more effective PDE4B inhibitor than Roflumilast.

**Figure 5A.**
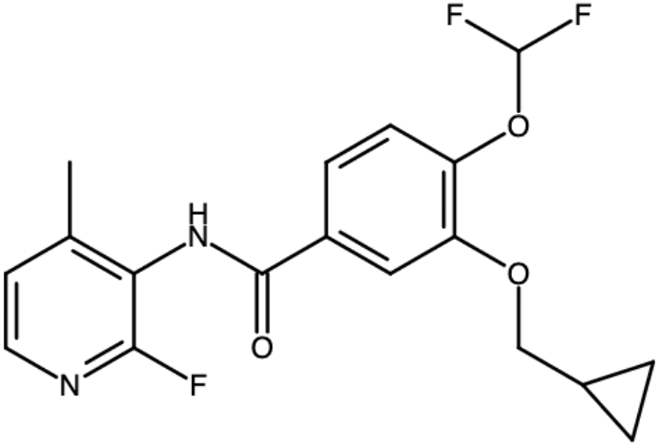
2D structure of Candidate 1.

**Figure 5B.**
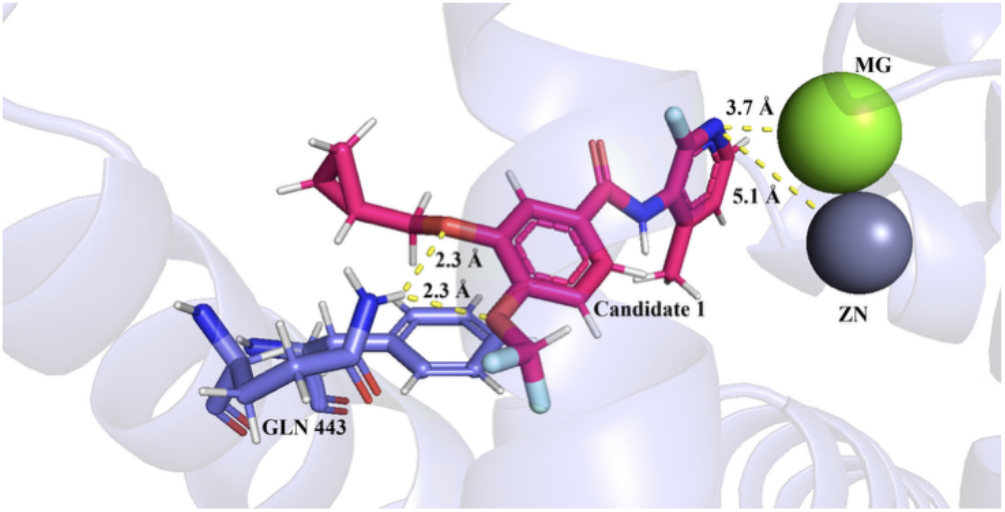
Dialkoxyphenyl (pink) bidentate hydrogen bonding with Q443 (blue), and pyridine interacting with metal ions (green and grey).

**Figure 5C.**
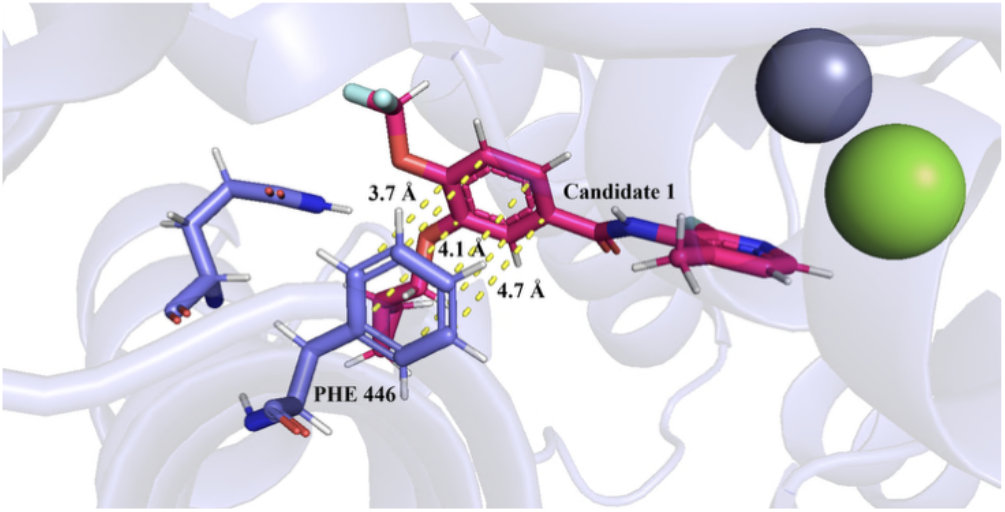
Dialkoxyphenyl (pink) Pi-Pi stacking with F446 (blue).

**Table 2.** Docking score and ADMET properties of Candidate 1.

| Name | Docking Score | Molecular Weight (g/mol) | H-bond Acceptors | H-bond Donors | XLogP |
| --- | --- | --- | --- | --- | --- |
| Candidate 1 | -12.61 | 366.33 | 5 | 1 | 3.81 |
| Roflumilast | -11.50 | 403.21 | 5 | 1 | 4.23 |

### Chemical Intuition Driven Design of Candidate 2

Following the computationally driven design of Candidate 1, the design of Candidate 2 (Figure 6A) followed the hypothesis by chemical intuition that more modifications can be made to the pyridine moiety to further improve on the metal ion interaction in effort to improve on binding affinity. Chemical intuition for a second analog, Candidate 2, builds on the bioisosteric analog established for Candidate 1. Modifications to the pyridine ring were experimented to maximize interactions with the PDE4B active site.

Initial hypothesis focused on building the electron density of the pyridine moiety toward the metal ion pocket to strengthen this interaction. Inspired by the metal-coordinating carbonyl group in Zardaverine, also a PDE4 inhibitor, the addition of a carbonyl group was first considered.^5^ However, introducing a carbonyl at this site would disrupt the aromaticity of the pyridine ring. This was an undesirable outcome, as aromaticity is important for preserving metal ion interactions. As a result, a single alcohol group was introduced ortho to the nitrogen of the pyridine ring and opposite the fluorine substituent. In addition to providing more electron density to the pyridine moiety and maintaining aromaticity, the alcohol group also provided additional hydrogen bond donors and acceptors. This chemically intuitive modification was observed to improve binding affinity, supported by active site interactions.

With this chemical intuition in mind, the bioisosteric analog of Candidate 2 was built and computational studies were performed. It had a docking score of −15.31, a 33% improvement compared to Roflumilast and a 21% improvement compared to Candidate 1 (Table 3). The improvement in the docking score is due to the observed formation of a new hydrogen bond between the added alcohol group on the pyridine ring and the backbone of residue T345, observed at 2.2 Å (Figure 6D). Notably, this hydrogen bond is formed on the opposite side of the ligand from the dialkoxyphenyl group where it forms its bidentate hydrogen bonds, resulting in larger surface area of stabilization when interacting with the active site.

**Table 3.** Docking score and ADMET properties of Candidate 2.

| Name | Docking Score | Molecular Weight (g/mol) | H-bond Acceptors | H-bond Donors | XLogP |
| --- | --- | --- | --- | --- | --- |
| Candidate 2 | -15.31 | 382.33 | 6 | 2 | 3.3 |
| Roflumilast | -11.50 | 403.21 | 5 | 1 | 4.23 |

Additional improvements in active site interaction are seen other than the new hydrogen bond, such as the introduction of the alcohol group further improves on the strength of the metal ion interaction. The additional electron density from the alcohol group further strengthens the metal ions interactions, while maintaining the dialkoxyphenyl pharmacophore and pyridine aromatic ring of the bioisosteric analog of Candidate 1. The dialkoxyphenyl group maintains its bidenate hydrogen bond with Q443 at 2.3 Å and 2.4 Å, and displaced Pi-Pi stacking interaction with F446 at about 4.5 Å (Figure 6B and 6C). The pyridine ring maintains its interaction with the metal ion pocket (Figure 6B), and the new observed hydrogen bond with T345. With additional supporting computational evidence from the new hydrogen bond formation and improvement in docking score, Candidate 2 shows strong binding affinity to the PDE4B active site and shows potential to be the lead novel drug candidate.

The ADMET properties of Candidate 2 also had to be evaluated (Table 3). As with Candidate 1, no violations of Lipinski’s Rules or toxicophores were detected, which supports Candidate 2 to be safely administered as oral treatment. The molecular weight increased to 382.33 g/mol due to the added alcohol group, as expected. Importantly, Candidate 2 had the lowest calculated XLogP value of 3.3 among all analogs, further supporting good oral bioavailability. Computational efforts showed that Candidate 2 had the move improved docking score, maintained essential active site interaction with an improvement in the addition of a hydrogen bond, and favorable ADMET properties, Candidate 2 shows the most potential to be the best novel drug candidate compared to Candidate 1 and Roflumilast.

**Figure 6A.**
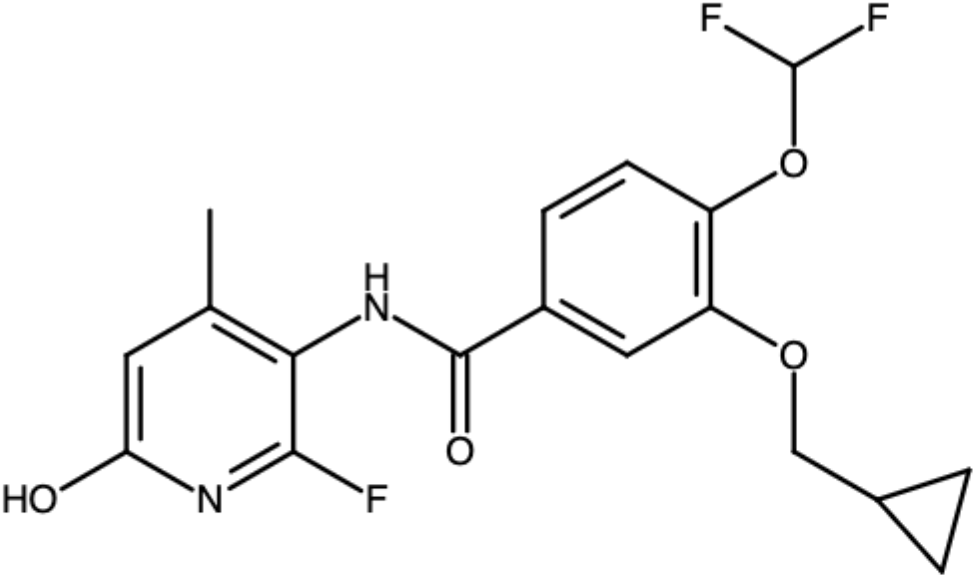
2D structure of Candidate 2.

**Figure 6B.**
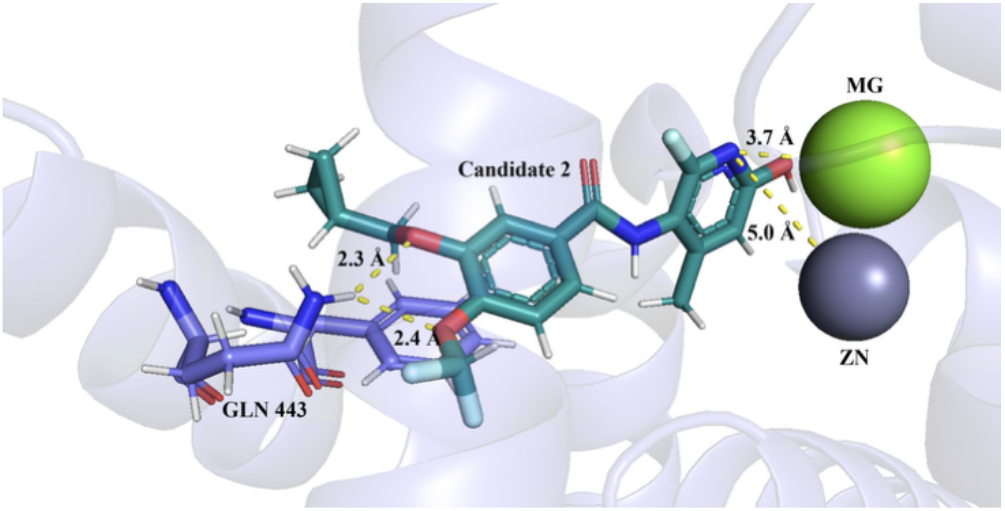
Dialkoxyphenyl (cyan) bidentate hydrogen bonding with Q443 (blue), and pyridine interacting with metal ions (green and grey).

**Figure 6C.**
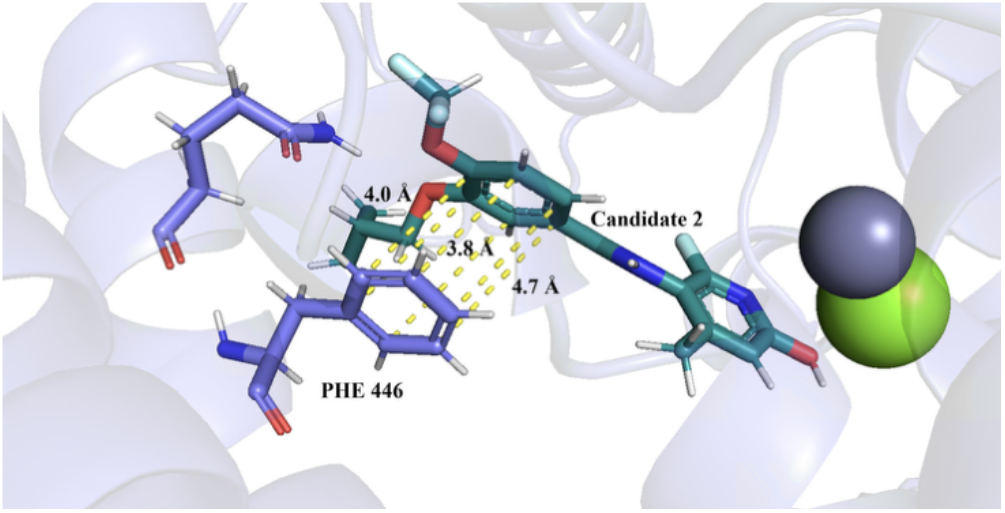
Dialkoxyphenyl (cyan) Pi-Pi stacking with F446 (blue).

**Figure 6D.**
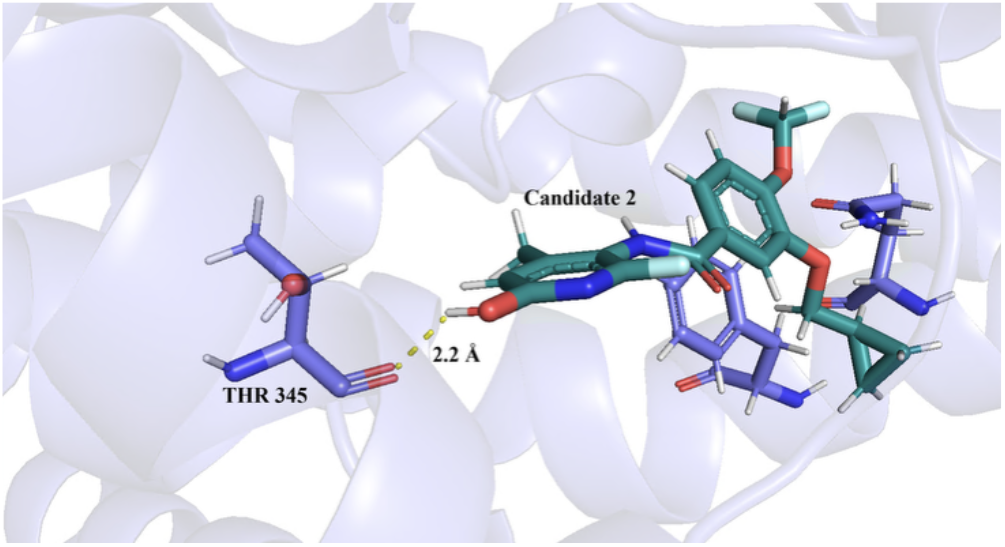
Hydrogen bond between additional alcohol group (cyan) and T345 (blue).

### Homology Analysis

Initial preclinical animal studies must be done to establish safety and efficacy first and foremost, and potential off-target effects in beneficial and pathogenic organisms following. To identify appropriate animal models that will give the most accurate model interaction of the novel candidate drugs with its protein, homologous proteins of the model organisms must be first identified. In efforts to achieve this, a homology analysis of the PDE4B protein was performed across 4 model animal organisms, as well as two homology analyses of a handful of microbes to screen for beneficial or pathogenic off-target effects. This gives basis for sequence alignment and analysis, which will guide the hypotheses for model animals for preclinical studies and off-target effects in microbes.

To establish which animal models were suitable for PDE4B, BLAST was used to identify homologous proteins across several species of organisms.^27^ Mus musculus (mice) had an average of 97% query cover and percentage identity average of 89%, while three Macaca species (Macaca fascicularis, Macaca mulatta, and Macaca nemestrina) (macaques) had an average of 94% query cover and percentage identity average of 84%. These high values indicate strong evolutionary conservation across most homologous proteins of the organisms and suggest both species are appropriate models for preclinical testing. In contrast, for Oryctolagus cuniculus (rabbits), conservation dropped drastically after the top 15 homologous proteins, indicating less similarity in the protein sequence and less suitable as a model organism for these novel drug candidates. Finally, Beagle (ID: 9615) was not available for search in the BLAST database, resulting in the inability to proceed with homology analysis for Beagle. Preliminary BLAST search for homologous proteins was essential to ensure that the model organisms being looked into will actually have observable interactions with the novel drug candidates.

Following homologous protein search, the active site residues essential for binding affinity had to be analyzed to check for conservation across the model organisms. Sequence alignments performed in Jalview revealed that essential residues, such as Q443 and F446 in the Q pocket, and T345, were fully conserved in both mice and macaques.^28^ Conservation of these residues supports the hypothesis that both novel drug candidates will maintain active site interactions needed for high binding affinity to PDE4B. Conservation of these residues support the bidentate hydrogen bonding and stabilization of the dialkoxyphenyl structure, and alcohol hydrogen bonding in the active site of PDE4B in these organisms. In conclusion, the conservation of the residues, ADMET properties and affinities should be similar to human homologous proteins when using mice and macaques as model animals in preclinical trial studies. While homology analysis strongly supports the hypothesis of mice and macaques as animal models, it is important to note the limitations utilizing computational studies. Actual outcomes can vary, thus can only be guaranteed through empirical studies.

To hypothesize the possibility of off-target effects on beneficial microbes, BLAST searches were performed to identify homologous proteins of PDE4B for common beneficial microbes including Lactobacillus, Bifidobacterium, and Akkermansia muciniphila.^27^ No significant PDE4B homologs were identified in any of these species, suggesting a low probability that the novel drugs would interact with the beneficial microbes. Consequently, this suggests that patients treated with either novel drug candidate are unlikely to experience off-target effects due to drug interaction with these beneficial microbes. Homology analysis results support the hypothesis that patients should be safe in regard to novel drugs potentially interacting with beneficial organisms.

The hypothesis of possibility of off-target interactions on pathogenic microbes is also notable for drug safety and efficacy. BLAST search concluded that Staphylococcus aureus, Haemophilus influenzae, and Tinea revealed no significant PDE4B homologs.^27^ However, in Candida albicans, a single homologous protein was found with 68% query coverage and 25.8% sequence identity. Although these values are low, there is a possibility of interactions between the novel drug candidates and the homologous proteins in Candida albicans. In hypothesizing the potential off-target effects of novel drugs from microbes, it is important to consider both presence and absence of any amount of protein when performing a homology analysis. Further sequence alignment analysis is required to identify conservation of residues and specific active site interactions.

Following the positive search results of pathogenic microbes, sequence alignment of the Candida albicans homologous protein confirmed that the three key active site residues (Q443, F446, and T345) are conserved. This suggests a high possibility of binding of the two novel drugs to the organism’s homologous protein active site, which supports the possibility of drug sequestration or off-target effects due to wrong host inhibition. In a clinical context, such interactions might reduce drug potency or cause unintended side effects if Candida albicans is present in a patient. While homologous analysis and sequence alignment provides substantial support for this hypothesis for off-target effects of pathogenic microbes, empirical studies have to be executed in order to fully guarantee or validate any computational study.

Though homology analysis is a great resource and provides a starting point for understanding how novel drugs can affect patients directly or indirectly, clinical studies have to follow to ensure safety and efficacy, and provide true empirical evidence. Provided that declaration, two animal models (mice and macaques) were presented to be suitable for preclinical animal studies, and the pathogenic microbe (Candida albicans) shows possibility of causing off-target effects if present in patients undergoing treatment of the novel drugs proposed.

## CONCLUSION

Plaque psoriasis, the most prevalent form of this chronic skin condition, manifesting in itchy and patchy skin which ultimately drains patients’ quality of life if proper treatment is not sought. Despite efforts to combat this disease, with no cure and limited options of current treatments leads to the collective goal of progressive innovation for drug development and discovery.

In effort to be a part of the collective goal, we computationally designed, alongside chemical intuition, and evaluated two novel PDE4B inhibitor drug candidates in this study, both structurally derived from Roflumilast. Both candidates had observed improvement in docking scores and more favorable ADMET properties for good oral bioavailability compared to Roflumilast. These improvements are largely attributed to intentional modifications of functional groups of analogs to tailor towards more favorable active site interactions. Out of all bioisosteric analogs evaluated, Candidate 1 and Candidate 2 have the most promising potential as the next novel drug based on the the improvement in docking scores and XLogP value, as well as ADMET properties and suggests significant binding affinity.

While computational studies were the basis of this research study, given almost all the evidence providing support for proposed hypotheses, it is still data that was generated from statistical computational software. Though evaluation was based on human judgement, there is room left for significant computational bias and error. Observations and conclusions, though highly valuable and provide important preliminary information, are limited and empirical studies must be performed to guarantee drug interactions with active site, safety and efficacy, and off-target effects. This research study then presents an essential next step if the proposed novel drugs were to be moved onto the next stage in development.

To support the next steps outside of computational studies into preclinical studies, comprehensive homology analyses and sequence alignments were conducted to understand animal models ideal for clinical studies, as well as what off-target effects are to be expected. These analyses identified mice (Mus musculus) and three macaque species (Macaca fascicularis, Macaca mulatta, and Macaca nemestrina) as suitable animal models for future preclinical animal studies, based on high conservation of residues in the homologous PDE4B protein in the organisms researched. Simultaneously, the analysis of conservation of the same active site residues of homologous protein PDE4B in Candida albicans presents a challenge of off-target effects. This further supports the need for studies that evaluate the general safety and efficacy of the proposed novel drug candidates in vivo.

With that said, the two proposed drug candidates show promising potential to be the next novel drug candidates of PDE4B inhibition, in light of combating the chronic autoimmune disease, plaque psoriasis.

## ACKNOWLEDGEMENT

This work was supported by the University of California Davis, the National Science Foundation (award nos. # 2315767, 2118138 and 1827246) and BioMADE FA8650-21-2-5028. The content is solely the responsibility of the authors and does not necessarily represent the official views of the National Science Foundation, other sponsors or UC Davis.

